# Chromosome-level *de novo* assembly of the pig-tailed macaque genome using linked-read sequencing and HiC proximity scaffolding

**DOI:** 10.1101/635045

**Authors:** Morteza Roodgar, Afshin Babveyh, Lan Huong, Wenyu Zhou, Rahul Sinha, Hayan Lee, John B. Hanks, Mohan Avula, Lihua Jiang, Hoyong Lee, Giltae Song, Hassan Chaib, Irv L. Weissman, Serafim Batzoglou, Susan Holmes, David G. Smith, Joseph L. Mankowski, Stefan Prost, Michael P. Snyder

**Affiliations:** Department of Genetics, Stanford University, Stanford, California, 94305, USA; Institute for Stem Cell Biology and Regenerative Medicine, Stanford University School of Medicine, Stanford, CA, USA; Institute for computational and Mathematical Engineering, Stanford University, Stanford, CA, 94305, USA; Center for Genomics and Personalized Medicine, Stanford University, Stanford, California, 94305, USA; Stanford Research Computing Center, Stanford University, Stanford, California, 94305, USA; School of Computer Science and Engineering, Pusan National University, Busan 46241, South Korea; Department of Computer Science, Stanford University, Stanford, CA, USA; Department of Statistics, Stanford University, Stanford, CA, 94305, USA; California National Primate Research Center, University of California, Davis, CA, 95616, USA; Department of Molecular and Comparative Pathobiology, Johns Hopkins University School of Medicine, Baltimore, MD, 21205, USA; LOEWE-Centre for Translational Biodiversity Genomics, Senckenberg, Frankfurt am Main, Germany; South African National Biodiversity Institute, National Zoological Garden, Pretoria, South Africa

## Abstract

Old world monkey species share over 93% genome homology with humans and develop many disease phenotypes similar to those of humans, making them highly valuable animal models for the study of numerous human diseases. However, the quality of genome assembly and annotation for old world monkeys including macaque species lags behind the human genome effort. To close this gap and enhance functional genomics approaches, we employed a combination of *de novo* linked-read assembly and scaffolding using proximity ligation assay (HiC) to assemble the pig-tailed macaque (*Macaca nemestrina*) genome. This combinatorial method yielded large scaffolds at chromosome-level with a scaffold N50 of 127.5 Mb; the 23 largest scaffolds covered 90% of the entire genome. This assembly revealed large-scale rearrangements between pig-tailed macaque chromosomes 7,12, and13 and human chromosomes 2, 14, and 15.

## Introduction

Old world monkeys including macaques share approximately 93% homology with the human genome, and therefore are valuable model organisms for the study of human infectious and genetic diseases (1, 2). Pig-tailed macaques (*M. nemestrina*) have been widely used as animal models for human infectious diseases, such as human immunodeficiency virus (HIV) and Simian immunodeficiency virus (SIV) (3-8) infections, and noninfectious diseases (e.g., neurodegenerative diseases) (9, 10).

Different species of nonhuman primates (NHP) exhibit varying levels of susceptibility to diseases. Studying the variation in disease susceptibility among primate species (including humans) requires reliable reference genomes that enable cross-species comparative genomics (5). However, the qualities of the genome assemblies and annotations for some of the old world monkeys are inferior to that of the human genome (11). A major limitation of the current pig-tailed macaque genome assembly (Mnem_1.0) is its reliance on assembly methods that do not yield large continuous genome scaffolds. This hinders comparative genomics studies of disease-related features that might vary, both among NHP species and between these species and humans.

Here we present a high-quality chromosome-level assembly of the pig-tailed macaque (*M. nemestrina*) genome by combining 10X Genomics-derived linked-reads based *de novo* assembly and scaffolding using proximity ligation (HiC) (Figure 1A) (12, 13). This high-quality assembly allowed us to identify large-scale structural variations compared to the human genome. In addition, we annotated the genome using RNAseq and proteomics data from induced pluripotent stem cells (iPSCs) lines derived from the peripheral blood mononuclear cells (PBMCs) of the same animal. Using this annotation, we inferred phylogenetic relationships among pig-tailed macaque (*M. nemestrina)*, rhesus macaque (*M. mulatta*), cynomolgus macaque (*M. fascicularis*), and human (*H. sapiens*).

**Figure 1.**
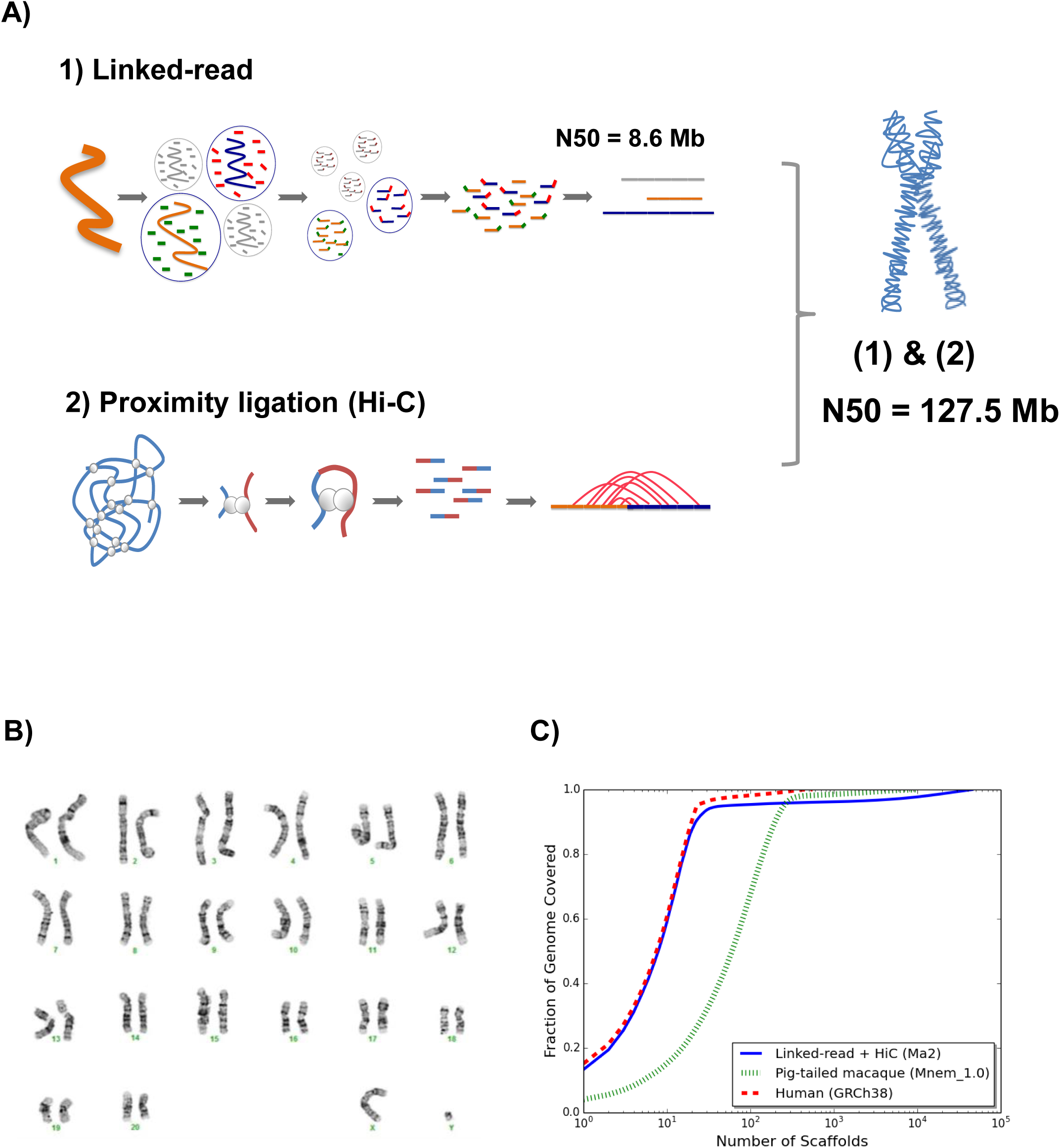
A) Schematic figure of the methods used for the assembly of the pig-tailed macaque genome (Ma2) 1) The Linked-read method resulted in scaffold N50 of 8.6Mb 2) A) Proximity ligation assay followed by scaffolding using HiRise method. This resulted in a scaffold N50 of 127.5 Mb (almost chromosome level). B) Karyotype of the pig-tailed macaque. C) Comparison of the number of scaffolds (X axis) and the proportion of the genome covered by the assembled scaffold (Y axis). The 23 largest scaffolds pig-tailed macaque (blue line) cover approximately 90 % of the genome. The red line presents the scaffold sizes of human genome (hg38) assembly and the green line is the current assembly of pig-tailed macaque on NCBI (Mnem_1.0).

## Results

### Chromosome-level assembly of the pig-tailed macaque genome

The presented *de novo* assembly (Ma2) represents a significant improvement in quality and scaffold size compared to the currently available assembly (Mnem_1.0) and is comparable in quality to the reference human genome (Figure 1C). Using a combination of linked-reads (10X Genomics Chromium System) and proximity ligation (HiC) based scaffolding we generated a genome assembly of a total length of 2.92 Gb with a scaffold N50 of 127.5 Mb. The 23 largest assembled scaffolds cover approximately 90 % of the entire pig-tailed macaque genome. We karyotyped the induced pluripotent stem cells (iPSCs) from the study animal. Since the pig-tailed macaque has 20 pairs of autosomes and a pair of sex chromosomes (Figure 1B)(14), each scaffold likely represents a single chromosome. With regard to scaffold sizes, the new pig-tailed macaque genome assembly (Ma2) is similar to that of the human genome which has been continuously improved over the last twenty years since its initial assembly (15). Using only the linked-reads method we obtained an assembly with scaffold N50 of 8.6Mb. However, using a combination of linked-reads and proximity ligation, we were able to increase the scaffold N50 to 127.5 Mb. Moreover, we observed significant improvements in reducing the extent of gaps in the assembled scaffolds. To evaluate the quality of our assembly, we ran Benchmarking Universal Single-Copy Orthologs (BUSCO) (16) using the OrthoDB mammalia database (Supplementary Figure1). We found 92% of complete BUSCO genes in the new pig-tailed macaque (Ma2) assembly, of which 89% were single-copy, 2.9% duplicated, 4% fragmented and 4.2% missing (Supplementary Table 1).

### Comparison of the new pig-tailed macaque genome with the human and rhesus macaque genomes reveals both extensive synteny conservation and genome re-arrangements

Pig-tailed macaques have 20 pairs of autosomes and one pair of sex chromosomes (Figure 1B)(14). Using the new genome assembly of the pig-tailed macaque, we performed synteny comparison of chromosomes between rhesus and pig-tailed macaque and human and pig-tailed macaque. Synteny analysis among the pig-tailed and rhesus macaque indicated a high level of homology between the two (Figure 2C). Synteny between human and pig-tailed macaque genomes showed large structural rearrangements such as a split of chromosome 7 of the pig-tailed macaque into chromosomes 14 and 15 in the human genome. In addition, chromosomes 12 and 13 of the pig-tailed macaque both align onto human chromosome 2 (Figure 2A). We further investigated the read pairing of the proximity ligation libraries for each of these chromosomes to validate the observed large structural rearrangements. The mapping of linked read HiC data on chromosomes 7, 12, and 13 of pig-tailed macaque supports the accuracy and reliability of the identified rearrangements (Figure 3A and 3B). To rule out biases from repeat homologies, we also conducted synteny analyses comparing the pig-tailed macaque genome with that of human and rhesus macaque after masking repeat elements (Figure 2A and 2B). This did not change any of the results discussed above.

**Figure 2.**
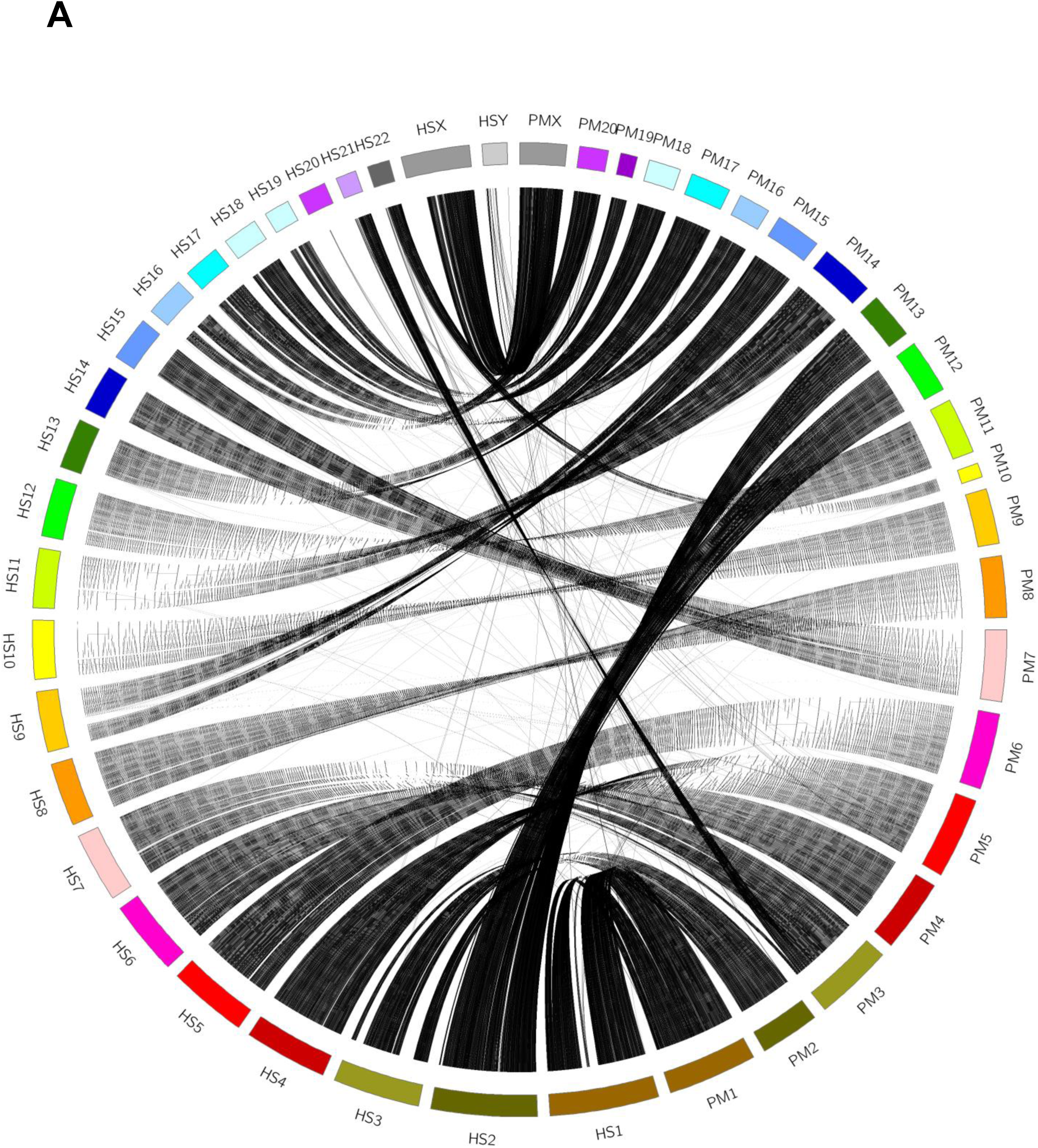
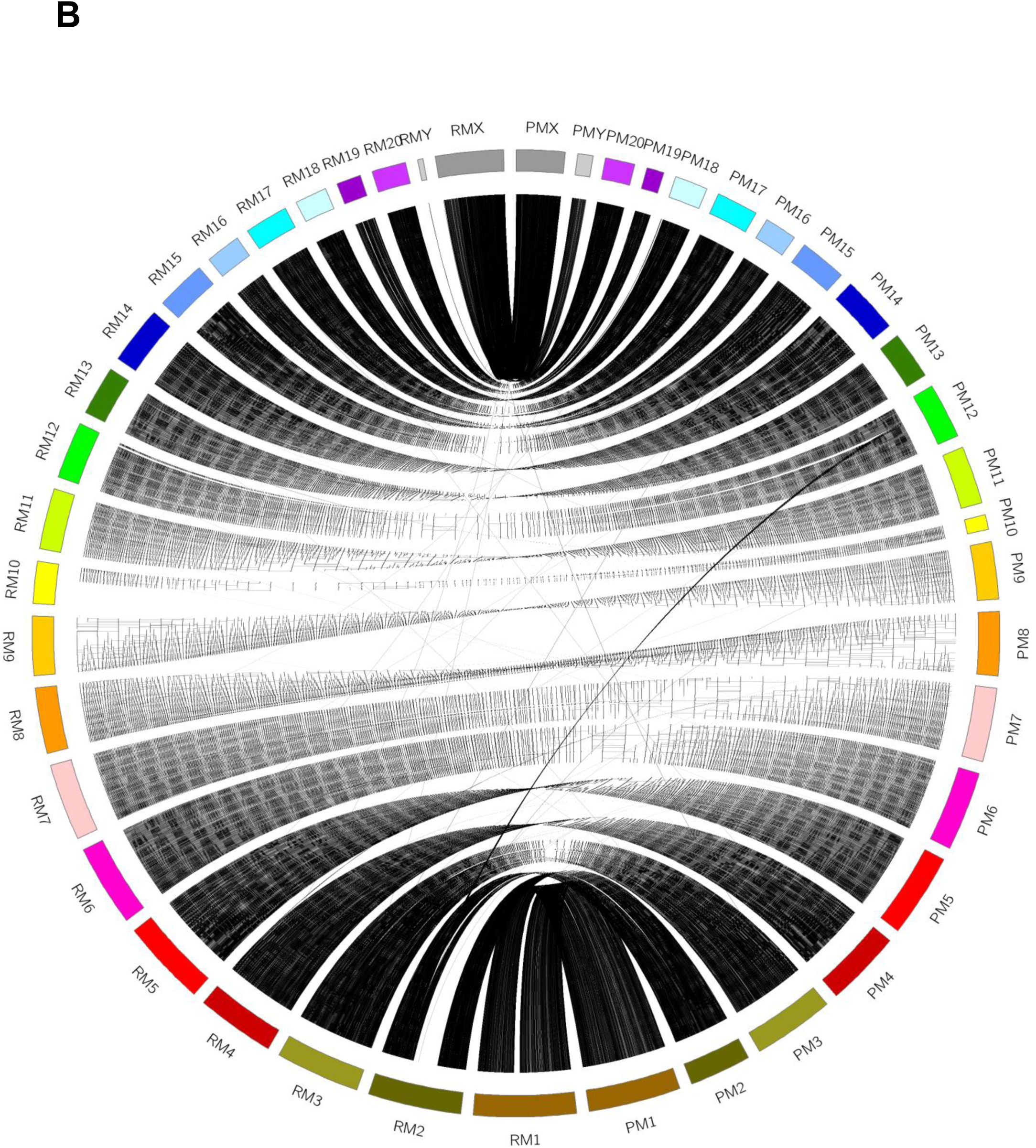
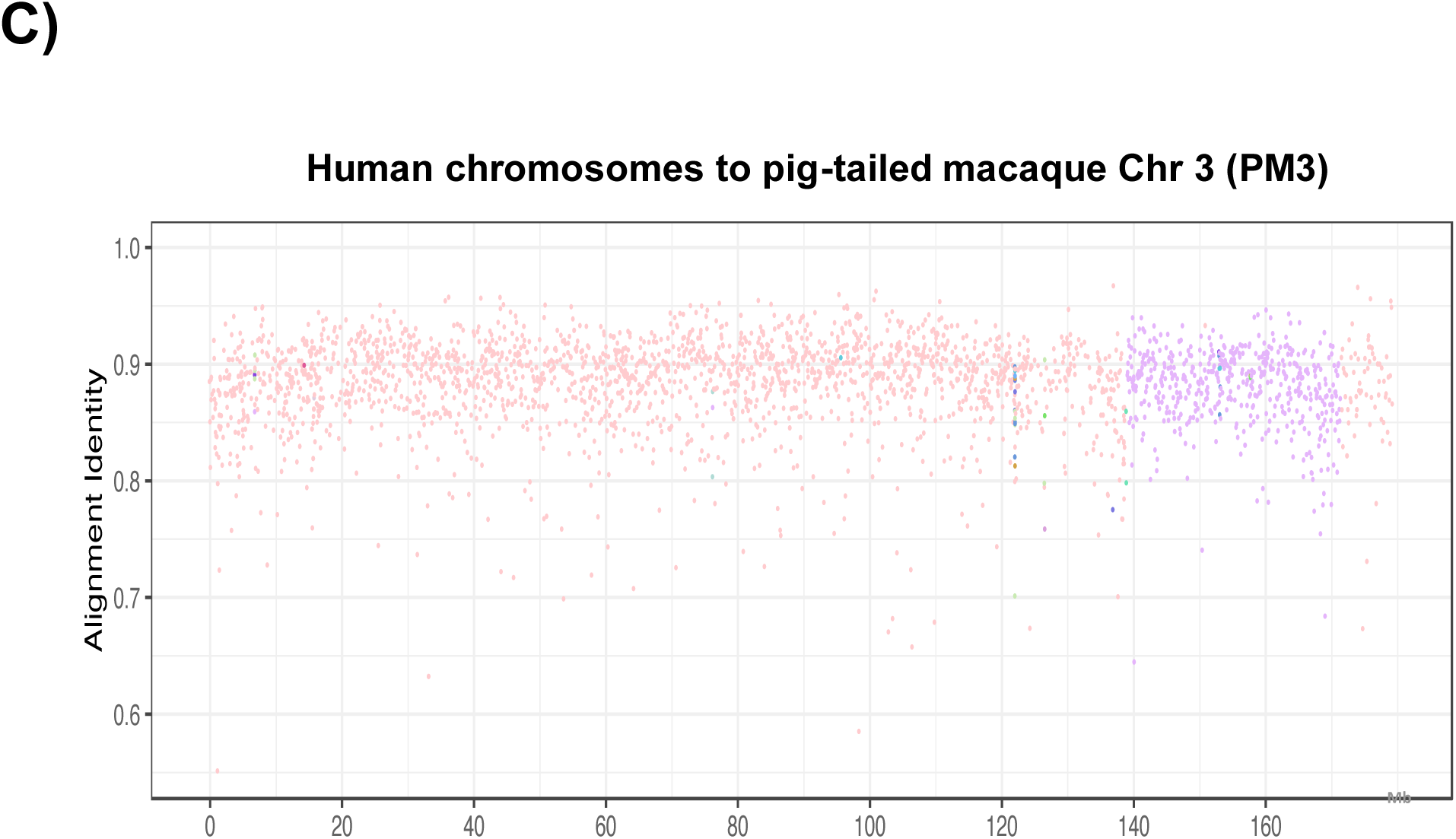
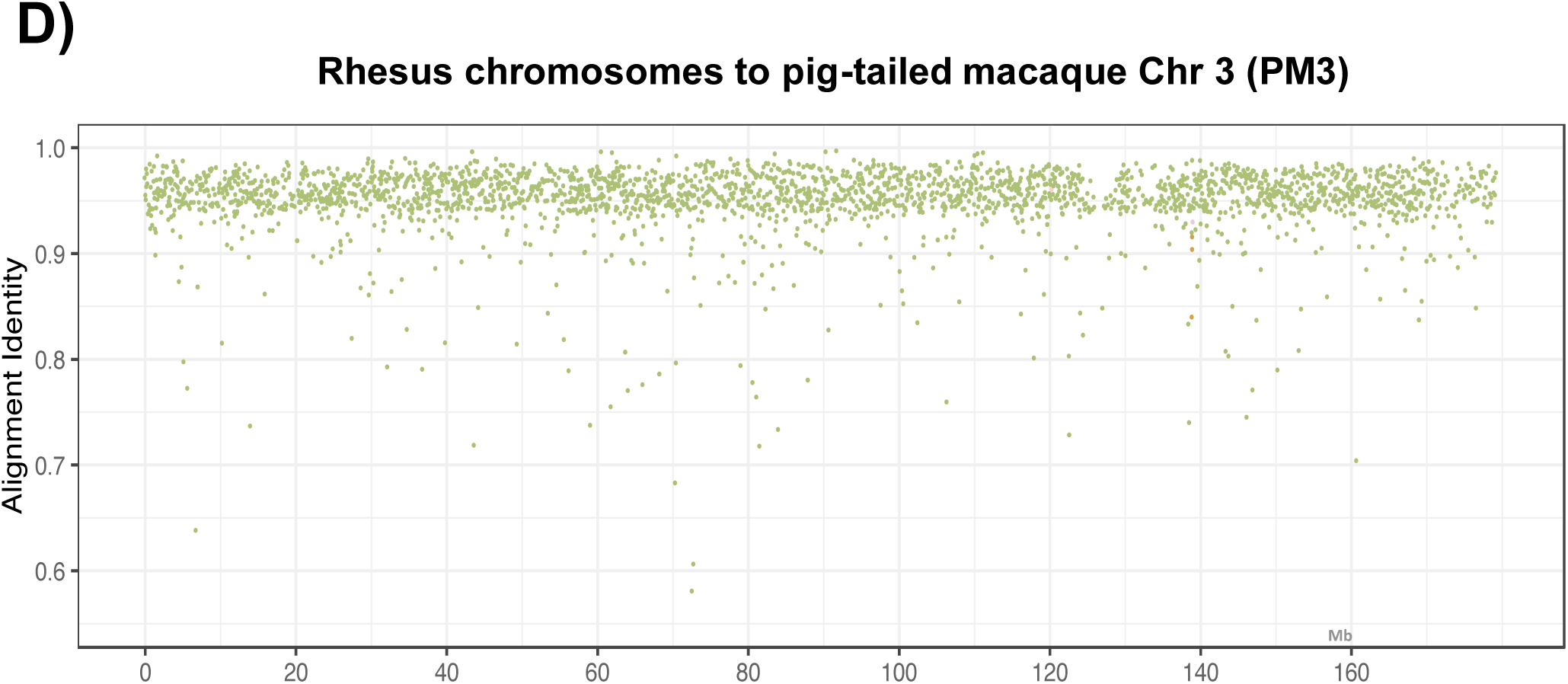
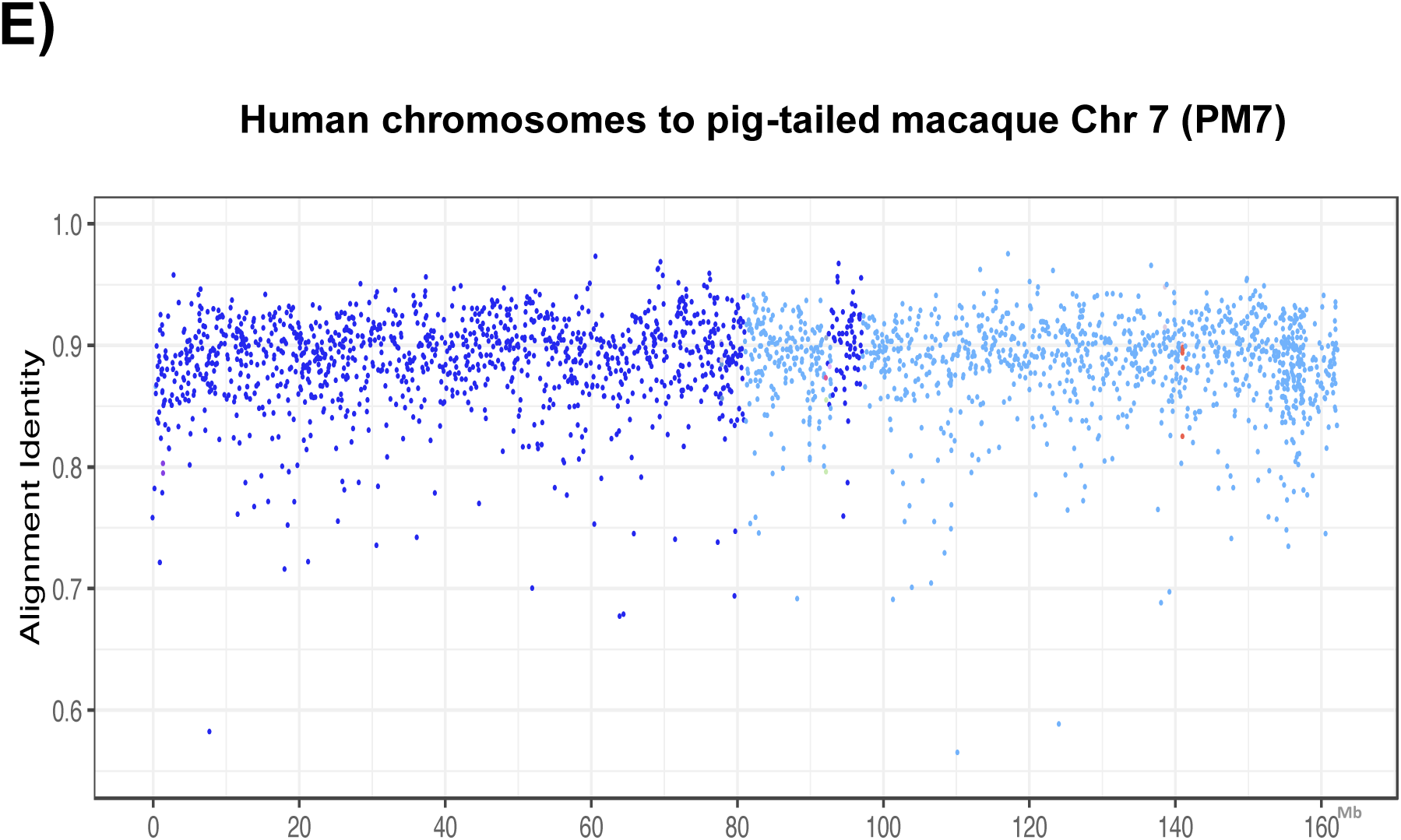
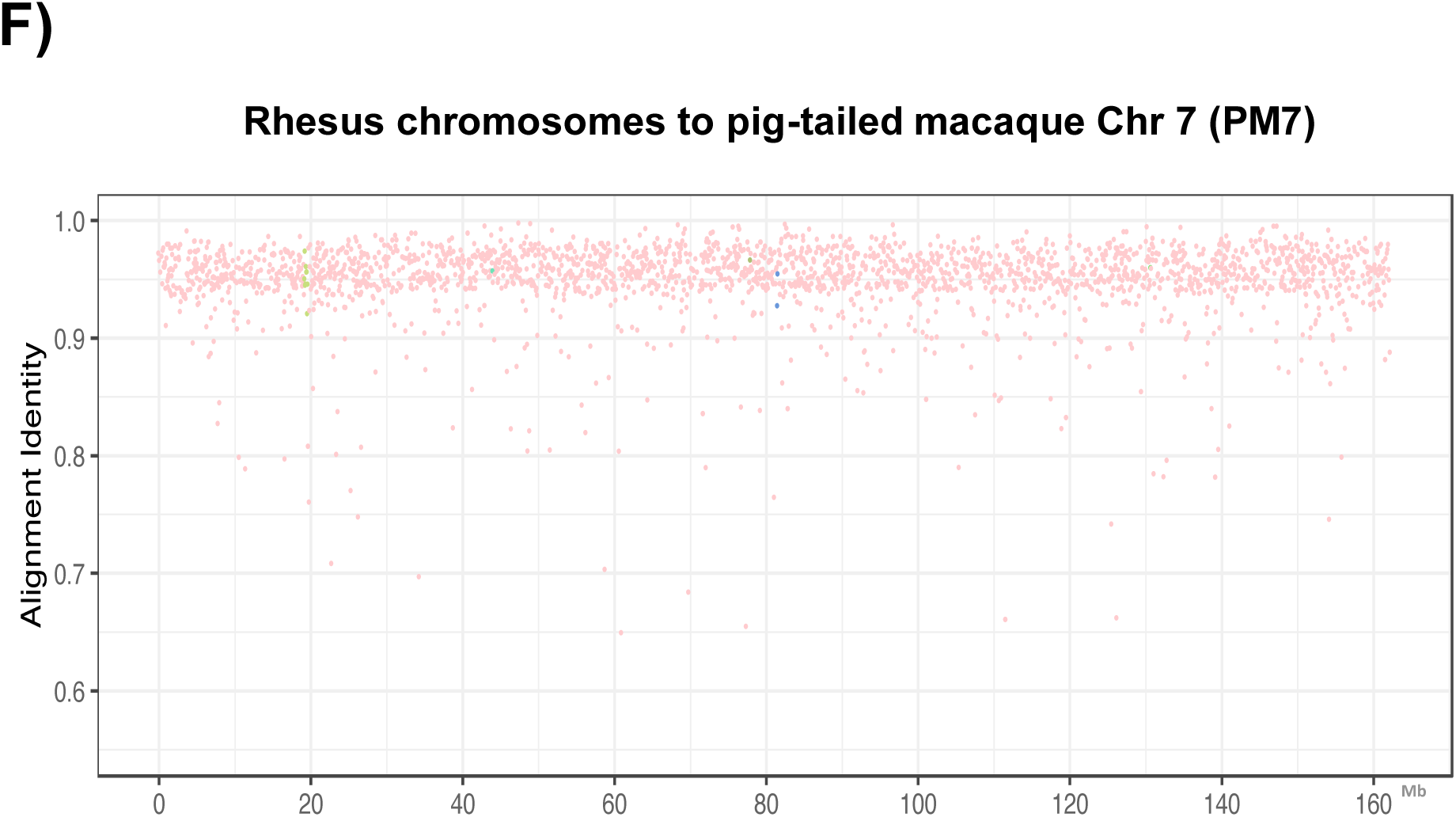
Synteny analysis and structural differences between pig-tailed macaque (PM) chromosomes 1 (PM1), PM chromosome 2 (PM2), through PMX and PMY with A) human chromosomes (HS1 through HSY), B) rhesus macaque (RM) chromosomes. C) Alignment identity score between human genome and pigtailed macaque chromosome 3 (PM3). D) Alignment identity score between rhesus macaque genome and pig-tailed macaque chromosome 3 (PM3). E) Alignment identity score between human genome and pigtailed macaque chromosome 7 (PM7) E) Alignment identity score between rhesus macaque genome and pigtailed macaque chromosome 3 (PM3). E) Alignment identity score between rhesus macaque genome and pigtailed macaque chromosome 7 (PM7).

**Figure 3.**
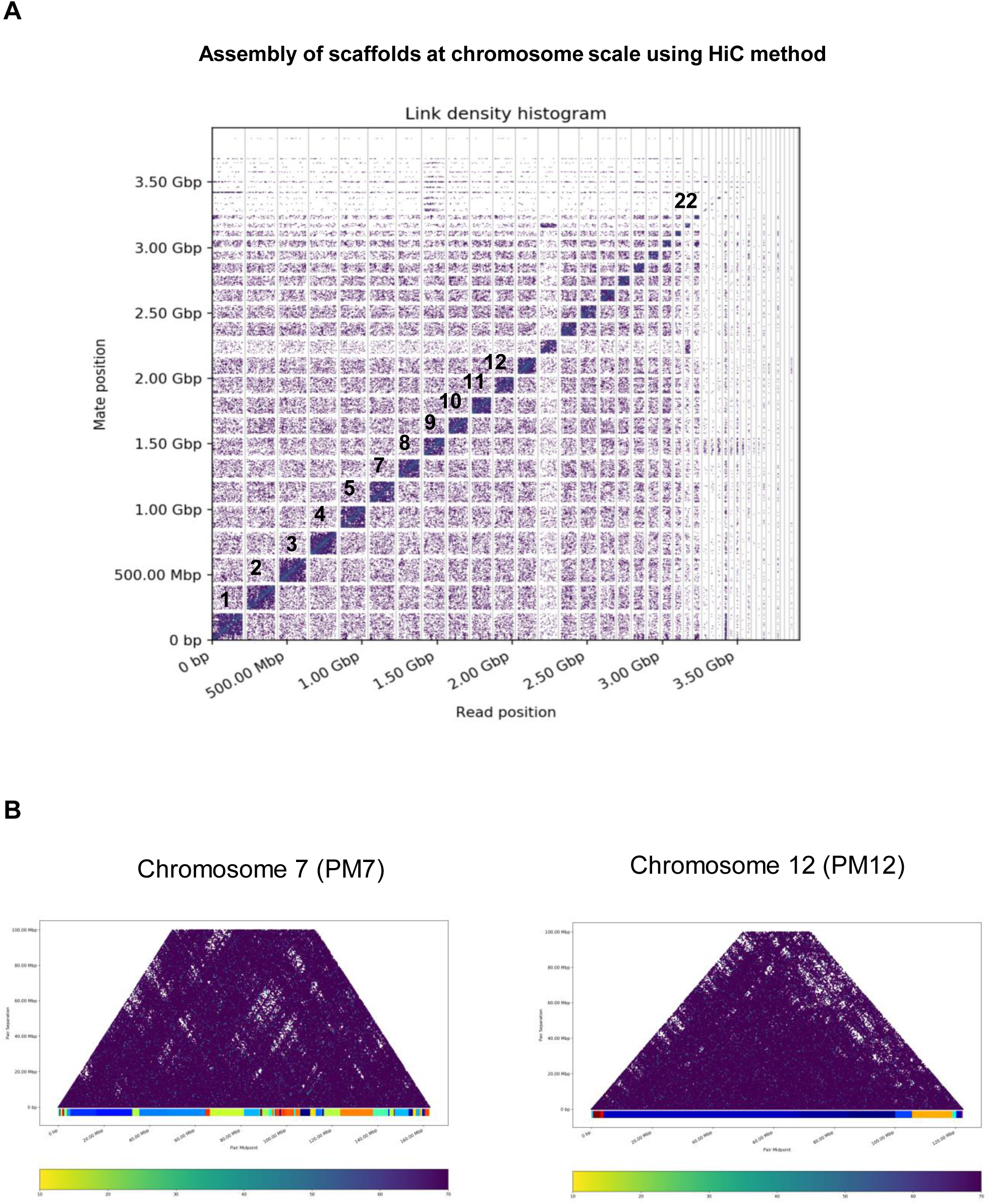
A) Linked density histogram of the assembled scaffolds of pig-tailed macaque genome. The number of dark boxes is the number of largest assembled scaffolds. B) Mapping of HiC read pairs on the pig-tailed macaque chromosome 7 (left) and the pig-tailed macaque chromosome 12 (right).

### Repeat Elements

Our annotation of repeat elements (Table 1) shows similar proportions to those present in rhesus and cynomolgus macaque, and human(1, 17). The assembly-wide repeat content fraction is 42%, which is also similar to that of other primates including humans (18).

**Table 1.** The statistics of repeats in the pig-tailed macaque genome.

|  | No. elements | Length occupied | Percentage of sequence |
| --- | --- | --- | --- |
| **SINEs:** | 1772888 | 371166008 bp | 12.70% |
| Alu/B1 | 1133107 | 290862183 bp | 9.95% |
| MIRs | 527205 | 73641539 bp | 2.52% |
| **LINEs:** | 982939 | 488123096 bp | 16.70% |
| LINE1 | 577434 | 382970388 bp | 13.10% |
| LINE2 | 349827 | 92659483 bp | 3.17% |
| L3/CR1 | 45394 | 9932162 bp | 0.34% |
| RTE | 9259 | 2409770 bp | 0.08% |
| **LTR elements:** | 594444 | 261853336 bp | 8.96% |
| ERVL | 112647 | 55029397 bp | 1.88% |
| ERV_L-MaLRs | 249607 | 105164349 bp | 3.60% |
| ERV_classI | 126831 | 79524160 bp | 2.72% |
| ERV_classII | 80815 | 15802293 bp | 0.54% |
| **DNA elements:** | 429884 | 102125714 bp | 3.49% |
| hAT-Charlie | 220542 | 44201979 bp | 1.51% |
| TcMar-Tigger | 101184 | 35948175 bp | 1.23% |
| **Unclassified:** | 54697 | 5632949 bp | 0.19% |
| **Total interspersed repeats:** |  | 1228901103 bp | 42.04% |

### Pig-tailed macaque (*Macaca nemestrina*) gene annotation

We annotated the pig-tailed macaque genome using RNAseq and proteomics data from an iPSC line generated from the same animal used for the genome assembly. The iPSC validation for this animal has been conducted as described in Roodgar et al. (2019). We also included orthologous proteins of closely related primate species for the annotation. This resulted in the identification of 39,579 transcripts. We then compared orthologous genes across the three macaque species and humans. We identified, 26,884 annotated pig-tailed macaque transcripts with no orthologs in either rhesus macaque or human, 8,490 transcripts with orthologs in both rhesus macaques and human, 2,153 transcripts with orthologs in humans only and 2,052 transcripts with orthologs in rhesus macaque only (Supplementary Figure S3A).

We next focused on comparing orthologous innate immune system transcripts among macaque species and human to determine whether differences in innate immune response orthologs existed between these species. We used a list of 844 innate immune system genes from the Innate DB database (19). Interestingly, we observed that several transcripts related to the immune system such as interleukin 2 receptor subunit alpha (IL2RA, RefSeq ID: NM_001032917.2), FAS (NM_001032933.2) (20-22) had orthologs in the three macaque species with no identified orthologs in humans. Among the list of innate immune genes, we also observed interleukin 31 (NM_001014336.1), interleukin L36 (NM_014440.2, NM_019618.3), and CD101 transcript variant 1 (NM_001256106.2) to have orthologs in human and pig-tailed macaque with no identified orthologs in rhesus or cynomolgus macaques. A list of the genes with orthologs in the three macaque species and no identified orthologs in humans is presented in Supplementary Table 2. We also observed that additional genes had annotated transcripts in the three macaque species with no identified ortholog in humans; these included AVPR1B (NM_001246222.1), the mutations of which correlate with anxiety, depression, and panic disorder in humans (23, 24).

### The use of proteomics data to improve the annotation

We conducted a two-step search to identify genes corresponding to each protein to improve the annotation of the pig-tailed macaque genome. The first search was against the initial pig-tailed macaque annotation database, created from the RNAseq data; in the second step, unmatched protein spectra were searched against the Human GENCODE19. Our initial Ma2 annotation database identified 9,031 proteins and the Human database identified extra 4,531 proteins. The total identified protein ID was 13,562 with FDR 1%. In a further attempt to improve the annotation, we added proteins GENCODE 28 database (22) to the input data used for the annotation using Maker2 (18). This resulted in a total number of 13,797 proteins used for the annotation. A list of all identified proteins can be found in Supplementary Table 3.

### Pairwise ortholog comparison of macaque species with human

We used POFF(25) to infer orthologous relationship between all the annotated genes in the three macaque species and humans. From the POFF graph-file output, we extracted the bit scores (a statistical value that represents sequence similarity) for each gene between human and each of the three macaque species. We aggregated the bit scores for each gene in the gene sets of interest (the list of innate immune genes and pluripotent stem cell genes) by computing the mean over all corresponding orthologs for each gene. For genes in the innate immune (IM) and pluripotent stem cell (PSC) gene set, the mean ortholog bit score difference to a corresponding human ortholog is similar for rhesus and cynomolgus macaques, exhibited by data points clustering along the 45 degree line (Supplementary Figure S3A&S3B). However, the mean bit score for several IM or PSC genes is higher for rhesus and cynomolgus compared to pig-tailed macaque, suggesting that the ortholog sequences of rhesus and cynomolgus macaques are more similar to human than those of the pig-tailed macaque. Some genes such as CXCL12 and EIF2S2 exhibit more similar sequences between humans and the pig-tailed macaque than between human and either rhesus or cynomolgus macaques (Supplementary Figure S3A&S3B).

### Reconstruction of the species phylogenetic tree

In this step, we applied POFF analysis to find orthologous sequences among five species: the three macaque species, human, and mouse. We observed that out of the total 124,011 human transcripts with an ortholog from at least one other species, 8,286 (6.7%) corresponded to innate immune genes. In contrast, among the total of 4,434 orthologs conserved in all five species, 905 (20.4%) belonged to the innate immune gene set. The over-representation of innate immune sequences in the conserved ortholog set (hypergeometric test p-value<1e-16) suggests that innate immune genes exhibit higher conservation than other genes. These immune genes include several cytokines, chemokines, and toll like receptors (TLRs).

We further reconstructed phylogenetic relationships among the three macaque species and humans using orthologous genes, using mouse as an outgroup. We reconstructed individual maximum likelihood-based gene trees for the 4,434 most conserved orthologs present in all the five species using Randomized Axelerated Maximum Likelihood (RAxML) (26). Out of the 4,434 orthologs identified in all five species, 2,339 (52.8%) follow the topology of phylogenetic tree 1 depicted in Figure 4A. We also investigated the proportion of the innate immune system genes (IM) or genes related to pluripotent stem cell genes (PSC) that follow each of the possible phylogenetic trees in the three macaque species, human and mouse (Figure 4A&B and Supplementary Figure S4). Our results indicate that the rhesus and cynomolgus macaques are closely related and exhibit a closer evolutionary distance to each other than either species exhibits to either human or pig-tailed macaque (Figure 4A). We observed that genes such as PDX1 and EIF2S2, follow the phylogenetic tree 6 in which the human and the pig-tailed macaque are closely related (Supplementary Figure S4).

**Figure 4.**
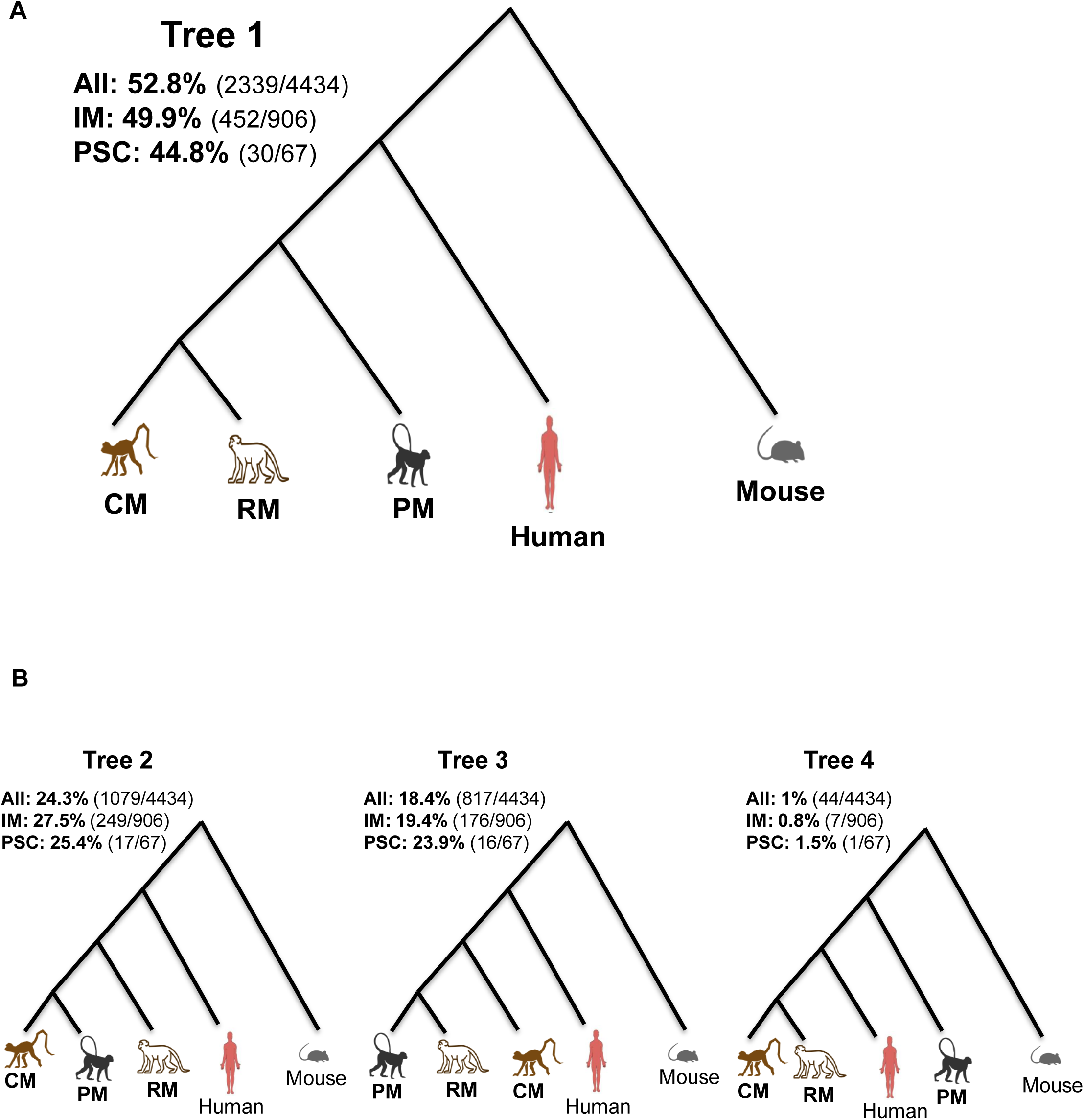
Reconstruction of the phylogenetic relationships among the three macaque species and human. A) The tree topology that is represented by most of the conserved genes among human, pig-tailed macaque (PM), rhesus macaque (RM), cynomolgus macaque (CM), and mouse as an outgroup. 52.8% of all the 4,434 most conserved orthologs follow the topology of tree 1. Also, 452 (49.9%) orthologs out of 906 most conserved innate immune genes (IM) and 30 (44.8%) out of 67 most conserved pluripotent stem cell genes (PSC) follow the topology of tree 1. B) Tree 2, 3 and 4 each representing different proportion of all of conserved orthologs, innate immune genes (IM) and pluripotent stem cell genes (PSC) among the three macaque species, human and mouse.

## Discussion

The pig-tailed macaque (*Macaca nemestrina*) is one of the species of old world monkeys widely used in biomedical research. However, identification of chromosomal synteny between the genomes of the pig-tailed macaque and humans has not been possible before. This limitation has been mainly due to the low continuity of the current genome assembly, in contrast to the high-quality assembly of the human genome. Recent advances in sequencing and library preparation methods allowed us to generate a genome assembly of chromosome-level quality. Here we present, to our knowledge, the first linked-reads and HiC-based assembly of a primate genome. In this study, we assembled the pig-tailed macaque (*Macaca nemestrina*) genome by combining two methods, linked-read *de novo* assembly and proximity ligation (HiC) based scaffolding. This method has resulted in a chromosome-level assembly of the pig-tailed macaque genome. The 23 largest scaffolds cover over 90% of the entire pig-tailed macaque genome, a result that is comparable to the human hg38 genome.

Synteny analysis between the human and pig-tailed macaque genomes revealed large scale chromosomal rearrangements in chromosome 14 and 15 of human with chromosome 7 of the pig-tailed macaque. Moreover, we observed synteny between chromosomes 13 and 14 of the pig-tailed macaque to human chromosome 2. Such large rearrangements were not observed between the pig-tailed and rhesus macaque chromosomes (Figure 2A&B). The proximity ligation assay data confirmed the authenticity of the observed rearrangements (Figure 3B).

Next, we reconstructed orthologous relationships among annotated genes of pig-tailed macaques, rhesus macaques, cynomolgus macaques, and human using mouse as an outgroup. Reconstruction of the phylogenetic tree confirms the closer evolutionary distance between rhesus and cynomolgus macaques than between humans and pig-tailed macaques (Figure 4A). The majority of the genes fit the species tree topology presented in Figure 4A. A genome-scale estimation of the coalescent-base species tree using ASTRAL (27) also confirmed the tree topology presented in Figure 4A (27). However, some genes do not follow the general consensus species tree (Figure 4B) (Supplementary Figure S4).

Because different species of NHP exhibit varying levels of susceptibility to diseases, understanding host responses such as the innate immune system is crucial for understanding the evolutionary changes that resulted in humans being susceptible to certain diseases. High-quality assemblies and annotations of NHP are essential for such cross-species comparisons. In this study we identified orthologs of some important innate immune response genes present in pig-tailed macaques and humans but absent in rhesus or cynomolgus macaques, some of which might play roles in disease susceptibility variation. We investigated the evolutionary tree topology for innate immune genes, because of the significant role of innate immune genes in diseases with similar pathophysiology in macaques and humans (e.g, SIV, HIV, and TB). However, the majority of the innate immune genes also follow the tree topology of the general consensus tree (Figure 4A). The comparison of the innate immune system orthologs is important to understand the variation in disease susceptibility among macaque species and humans. Comprehensive annotations of the three macaque species genomes are required for such comparative genomic studies.

Induced pluripotent stem cells (iPSCs) that can be derived from somatic cells (e.g., peripheral blood cells) have the potential to differentiate into various tissues. Here we used iPSCs of the pig-tailed macaque, generated from the PBMCs of the study animal, for the genome annotation. Differentiation of iPSCs to different lineages in future studies could be a tool for screening the functional roles of different genes and repeat elements that are important in early developmental stages and in disease modeling among primate species (28). Furthermore, we have investigated some genes with significant roles in pluripotent stem cell biology. The tree topology of KLF4 and c-Myc, two important genes in pluripotent stem cell biology (29), also followed the general consensus trees (Figure 4A). Pluripotent stem cell genes (PSC) are of particular interest because of adaptation of pig-tailed macaque iPSCs in maintaining pluripotency in human pluripotent culture media (Roodgar et al 2019). We also observed that human Wnt family member 8A (WNT8A) transcript variant 1 (NM_001300938) has an orthologous transcript in pig-tailed macaques but lacks an orthologous transcript in rhesus or cynomolgus macaques.

The presented assembly is of a single individual, while the human genome assembly increasingly represents some of the variations in highly variable regions across many individuals. There is a need for enhancing this assembly with a representation of pig-tailed macaque populations in the future

In conclusion, the chromosome-level assembly of the pig-tailed macaque genome provides a useful resource for the study of genome rearrangements, structural variation and their possible roles in disease susceptibility among primates. In addition, gene orthology relationship and reconstruction of phylogenetic trees among the three macaque species and human provide new insights into their evolutionary history. We investigated the evolutionary distance among the three macaque species and human with regards to the innate immune and pluripotent stem cell genes. Further investigation of evolutionary distances among these primate species and humans based on these and other gene sets can facilitate the selection of appropriate animal models for the study of different human diseases.

## Acknowledgements

This work used the Genome Sequencing Service Center by Stanford Center for Genomics and Personalized Medicine Sequencing Center, supported by the grant award NIH S10OD020141 and samples from the Johns Hopkins Pigtailed Macaque Breeding Colony supported by NIH U42OD013117.

## Materials and methods

### Study subject, sample preparation and DNA extraction

The study subject is a 14-year-old male pig-tailed macaque (*Macaca nemestrina*) from the pig-tailed macaque colony at Johns Hopkins University. Blood was drawn from the study animal following the Institutional Animal Care and Use Committee (IACUC) protocol at Johns Hopkins University. Peripheral blood mononuclear cells (PBMCs) were isolated from whole blood collected in ACD from the male pig-tailed macaque Ma2 by percoll gradient centrifugation. After separation, cells were stored viably in 90% FBS/ 10% DMSO in liquid nitrogen (30). Approximately one million PBMC were used for the extraction of High Molecular Weight (HMW) DNA using the MagAtrract HMW DNA Kit (Qiagen Inc., Valencia CA). The average size of the HMW DNA extracted DNA, approximately 55kb (Supplementary data), was measured using a Fragment Analyzer (Advanced Analytical Technologies, Inc., Ankeny, IA). Additionally, induced pluripotent stem cells (iPSCs) were derived from PBMCs as described elsewhere (Roodgar et al. 2019).

#### Chromosome karyotyping

The macaque-derived (*M. nemestrina*) induced pluripotent stem cell line derived from the study animal’s peripheral blood mononuclear cells (PBMCs) was harvested by standard cytogenetic methodology of mitotic arrest, hypotonic shock and fixation with 3:1 methanol-acetic acid. Chromosome slide preparations were stained by G-banding and classified by the standard *M. nemestrina* karyotype 1, 2. Analysis of 20 metaphase cells demonstrated an apparently normal male karyotype of 20 autosomes and two sex chromosomes (X and Y) in all cells (14, 31).

### Library preparation and sequencing

In order to generate a chromosome-level assembly, we prepared two different genome libraries.

#### 1) Linked-read library preparation

The 10X Genomics Genome Chromium platform (10X Genomics, Pleasanton CA) including Gel Bead-In-EMulsions (GEMs) generation and barcoding was used to generate the linked-read library (32). In summary, in a 10X Genomics Chromium microfluidic Genome Chip approximately 0.8-1.2ng of HMW DNA was combined with master mix and partitioning oil to create GEMs. Single HMW DNA molecules in each Gel Bead were barcoded with a unique 10X genomics barcoded primer. Upon dissolution of the Genome Gel Bead in the GEM, primers containing (a) an Illumina® R1 sequence (Read 1 sequencing primer), (b) a 16 nucleotide 10X Genomics Barcode, and (c) a 6-nucleotide random primer sequence were added into the fragments of DNA. The barcoded fragments of DNA ranging from a few to several hundred base pairs were pooled together when the GEMs were broken after incubation. Silane magnetic beads were used for the post GEM generation clean up to remove leftover biochemical reagents. Solid Phase Reversible Immobilization (SPRI) beads were used to optimize the appropriate DNA size range for library preparation (AMPure XP, BeckmanCoulter Inc, Indianapolis, IN, USA). Read 1 sequence and 10X Genomic barcodes were added during the GEM generation and P5, P7 primers, Read 2 and sample index were added during library construction after end-repair, A-tailing, and adaptor ligation and amplification. The constructed library was evaluated on a Bioanalyzer 2100 using Agilent High Sensitivity DNA (Agilent Inc., Santa Clara, CA, USA) and quantified using Kapa Library Quantification Kit (KappaBiosystems Inc., Wilmington, MA, USA). The sequencing of each sample was conducted on three lanes on Illumina Hi-seq 4000 sequencing platform generating approximately 1.8 billion of paired end read sequencing data with a 2×151bp read size (R1 and R2).

#### 2) HiC library preparation by Dovetail Genomics

Three Dovetail HiC libraries were prepared in as described previously (13). Briefly, for each library, chromatin was fixed in place with formaldehyde in the nucleus and then extracted. Fixed chromatin was digested with DpnII, the 5’ overhangs filled in with biotinylated nucleotides, and then free blunt ends were ligated. After ligation, crosslinks were reversed and the DNA purified from protein. Purified DNA was treated to remove biotin that was not internal to ligated fragments. The DNA was then sheared to a ∼350 bp mean fragment size, and sequencing libraries were generated using NEBNext Ultra enzymes and Illumina-compatible adapters. Biotin-containing fragments were isolated using streptavidin beads before PCR enrichment of each library. The libraries were sequenced on an Illumina Hi-seq 4000 sequencing platform (2×151bp). The number and length of read pairs produced for each library were 141 million for library 1; 88 million for library 2; 133 million for library 3. Together, these Dovetail HiC library reads provided 1990.23x physical coverage of the genome (1-50kb pairs).

### Genome assembly

Next, we assembled and scaffolded the data. The genome assembly was carried out using 10X Genomics’ Supernova2 assembler with default parameters, and the resulting assembly scaffolded using Dovetail Genomics’ HiRise scaffolder.

#### A. Linked-read library

First, we assembled the linked-reads using Supernova2 (10X Genomics Inc, Pleasanton, CA) with default parameters (14) on the Amazon Web Services (AWS) *x1.16xlarge* instance with *976 GB* computing memory and 2Tb of SSD disk and *i3.16xlarge* instance with *488 GB* computing memory and 2TB of NVMe disk. A system requirement for the genome assembly of an organism with a diploid genome size close to that of the human genome (∼3.2Gbp) requires approximately 284 GB of memory and 2TB of disk space for the data storage. The assembly quality was assessed using Benchmarking Universal Single-Copy Orthologs BUSCO (2) (Supplementary figure).

#### B. Scaffolding the assembly using HiRise

The input *de novo* assembly, shotgun reads, and Dovetail HiC library reads were used as input data for HiRise, a software pipeline designed specifically for using proximity ligation data to scaffold genome assemblies (4). Shotgun and Dovetail HiC library sequences were aligned to the draft input assembly using a modified SNAP read mapper (http://snap.cs.berkeley.edu). The separations of HiC read pairs mapped within draft scaffolds were analyzed by HiRise to produce a likelihood model for genomic distance between read pairs, and the model was used to identify and break putative misjoins, to score prospective joins, and make joins above a threshold. After scaffolding, shotgun sequences were used to close gaps between contigs.

##### Repeat identification

Using RepeatMasker (Smit, AFA, Hubley, R & Green, P. *RepeatMasker Open-4.0)*, we masked genomic interspersed repeats from the genome of the pig-tailed macaque. These repeats include simple repeats (1-5bases), tandem repeats, duplication of 100-200 bases that are more typically found at centromeres and telomeres of chromosomes, segmental duplications (large blocks of 10kb to 300kb), processed pseudogenes, retrotranscripts, SINES, LINES, DNA transposons, and retrovirus retrotransposons.

##### Synteny analysis

In order to conduct synteny analysis between pig-tailed macaques and humans and between pig-tailed and rhesus macaques, we first carried out pairwise whole-genome alignments using Satsuma Synteny (33). Next, we plotted the links using Circos(34).

##### Chromosome assignment to the 23 largest scaffolds

We took two approaches to assign scaffolds to chromosomes: 1) We used synteny analysis between pig-tailed and rhesus macaque to identify chromosomes of rhesus macaque genome that align to each of the 23 largest pig-tailed macaque scaffolds. 2) We blasted (11) validated genes with known loci on rhesus macaque genome onto the pig-tailed macaque genome to verify the assignments using the synteny information.

##### Annotation

RNAseq: mRNA was extracted from the iPSCs of the pig-tailed macaque using RNeasy Mini Kit extraction kit (Qiagen Inc. Valencia CA). Following cDNA synthesis, the RNAseq library was prepared using an Illumina TrueSeq RNA Library Prep kit V2 (Illumina Inc., San Diego CA) with unique dual index to avid index switching(35). Sequencing was conducted using a HiSeq4000 Illumina machine generating approximately 50 million paired reads (2×150bp, Illumina Inc., San Diego CA). The RNAseq data was subsequently used for annotation steps.

Gene annotation was then conducted using the software Maker2 (36). For the annotation of the genome we used RNAseq and proteomics data from the study animal’s PBMCs and iPSCs (generated from the PBMCs). To do so, RNAseq data was aligned to the genome using HiSat2 (37), and the assembly was conducted using StringTie (38). In addition to the RNAseq data from iPSC of the study animal, we added the pig-tailed macaque RNAseq and proteomics data available on the ENSEMBLE and NCBI data bases, as well as the proteomics data from human Genecode data base.

##### Proteomics Sample Preparation

Cells pellets were lysed in 100ul 6M GdmCl, 10mM TCEP, 40mM CAA and 100 mM Tris pH 8.5 buffer. Lysates were incubated at 95 °C for 5 min and briefly sonicated. Samples were boiled for 5min at 95°C and vortexed every 1min, then spinned 5mins + 10mins + 10mins, 12000g, taking supernatant for each step to continue. Protein concentration was measured using the BCA method. Trypsin (modified, from Promega, Madison, WI) was added at a protein to enzyme ratio of 50:1. Samples were incubated overnight at 37 °C. Peptide were added TFA to 1% then cleaned up using an Oasis HLB cartridge (1cc/10mg, Waters). Samples were dried by speed vac and dissolved in 100mM TEAB. Peptides were labeled with TMT 10plex reagent (Thermo Fisher) and combined at equal amounts, then vacuum spun till dry.

Waters 2D liquid chromatography (Waters MClass 2DnLC) was used for peptide separation by reverse phase chromatography at high pH in the first dimension followed by an orthogonal separation at low pH in the second dimension. In the first dimension, the mobile phases were buffer A (20mM ammonium formate at pH10) and buffer B (Acetonitrile). Peptides were separated on an Xbridge 300µm x 5 cm C18 5.0µm column (Waters) using 15 discontinuous step gradients at 2 µl/min. In the second dimension, peptides were loaded to an in-house packed 100µm ID/15µm tip ID x 28cm C18-AQ 1.8µm resin column with buffer A (0.1% formic acid in water). Peptides were separated with a linear gradient from 5% to 40% buffer B (0.1% formic acid in acetonitrile) at a flow rate of 300 nl/min in 112min, 170min and 180 min. The LC system was directly coupled in-line with an Orbitrap Fusion Lumos Tribrid Mass Spectrometer (Thermo Fisher Scientific) via a Thermo nanoelectrospray source. The machine was operated at 1.8-2.2 kV to optimize the nanospray with the ion transfer tube at 275 °C. The mass spectrometer was run in a data dependent mode. The full MS scan was acquired in the Orbitrap mass analyzer with resolution 120,000 at m/z 375-1500 followed by top 20 MS/MS in ion trap. For all sequencing events dynamic exclusion was enabled to fragment peptide once and excluded for 60s.

##### Proteomics data processing and analysis

The raw data were then processed with the software Proteome Discoverer (Thermofisher Scientific Inc.). Mass tolerance of 10ppm was used for precursor ion and 0.6 Dalton for fragment ions for the database search. The search included cysteine carbamidomethylation as a fixed modification. Acetylation at protein N-terminus and methionine oxidation were used as variable modifications. Up to two missed cleavages were allowed for trypsin digestion. Only unique peptides with a minimum length of 6 amino acids were considered for protein identification. The false discovery rate (FDR) was set as less than 1%.

##### Maker Proteomics for gene identification

We conducted a two step search. The first search was against our Ma2 annotated database; the unmatched spectra were searched against the Human Genecode19. Our Ma2 annotation database identified 9,031 proteins, and the human database identified an additional 4,531 proteins for a total of 13562 identified proteins with FDR 1%.

In a further attempt to improve the annotation using Maker2 (36), we added the Gencode 28 database (39) to the homology-based annotation step in order to increase the number of annotated genes on the pig-tailed macaque genome. After adding the Gencode genes to a new Maker run, the total number increased to 13,797 master proteins. This included 3,979 proteins from the Genecode human database and 9,818 proteins from the initial pig-tailed macaque Ma2 annotation. The Thermo Proteome Discoverer 2.1.0.81 software was used for protein mapping.

##### Pairwise bit score comparison

We applied POFF(25) to find orthologous sequences between the three macaque species and human, using mouse as an outgroup.

From the POFF output, we extracted the bit scores (a statistical value that represents sequence similarity) of each gene for their distance from human to each of the macaque species. We aggregated the bit scores for each gene in the gene sets of interest (list of innate immune genes and the list of pluripotent stem cell genes) by computing the mean over corresponding orthologs.

##### Reconstruction of phylogenetic relationships between the three macaque species and human

The POFF software (25) was used to detect orthology between gene sequences from the 5 species, pig-tailed macaque, rhesus macaque, cynomolgus macaque, human and mouse (40). The pig-tailed macaque sequences were derived from our own assembly and annotation of the genome. The mouse sequences were included for comparison and as an outgroup for rooting the subsequently estimated evolutionary trees.

In the next step, MAFFT (41, 42) was applied to individual ortholog groups, each containing five sequences corresponding to the species listed above. Aligned sequences were subsequently used as inputs for RAxML (26) to reconstruct the phylogenetic relationships between the five species. The Astral program (27) was then applied to generate a species tree from the individual gene trees. Next, the gene trees were rooted using mouse as outgroups. We reconstructed a total of15 possible phylogenetic tree topologies. We explored the assignment of all genes as well as the subset of immune genes (IM) and pluripotent stem cell (PSC) genes to the 15 tree topologies. This can provide information on which species are closer or further from human.

---

**Supplementary Table 1.**
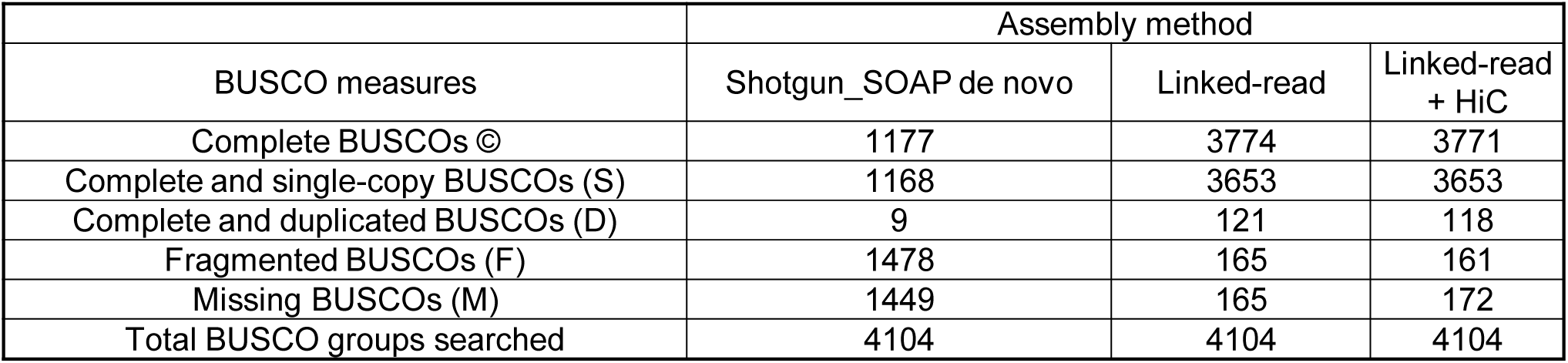
Benchmarking Universal Single-Copy Orthologs (BUSCO) for assembly evaluation.

**Supplementary Figure S2.**
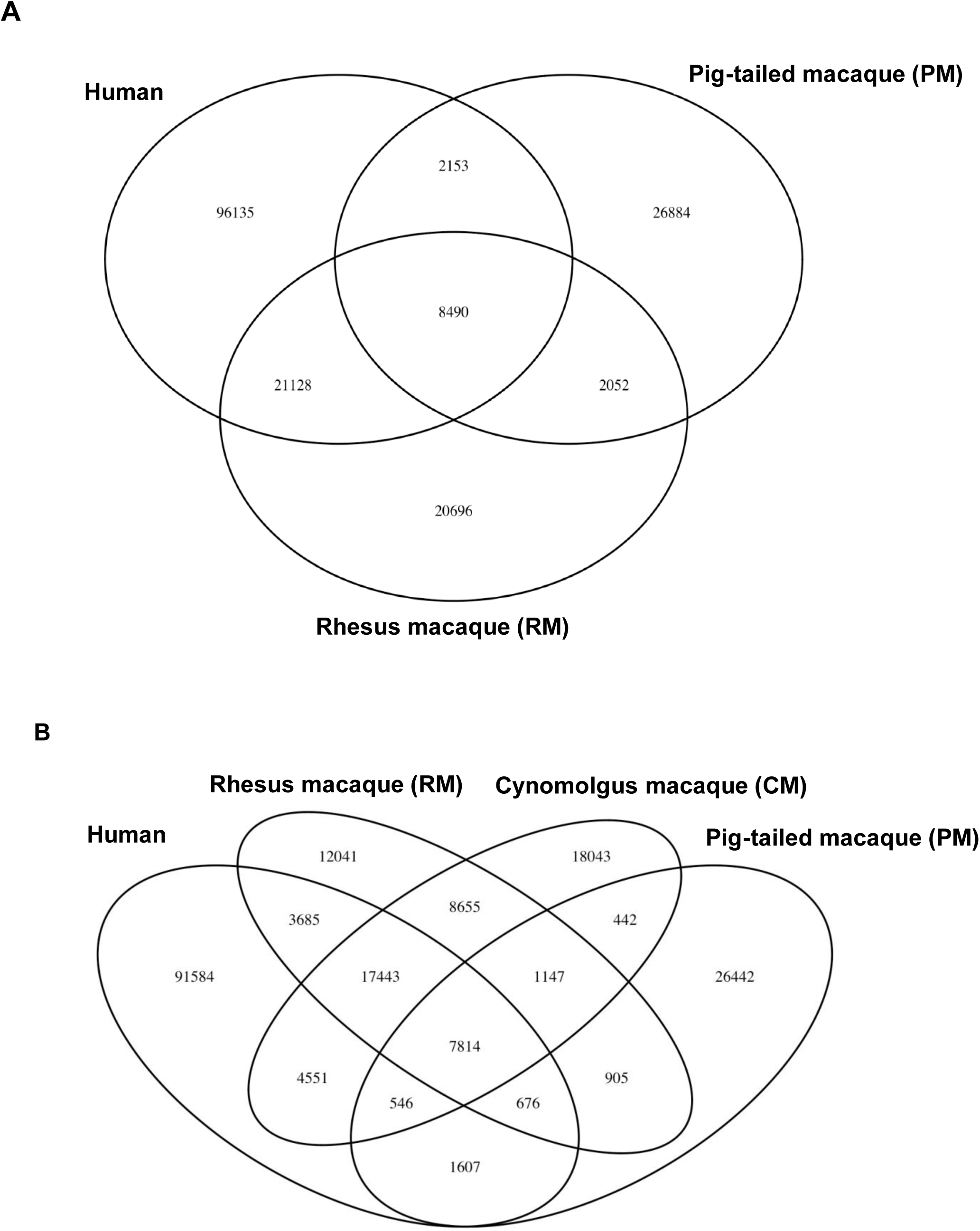
Venn diagram of the number of common orthologs A) among rhesus macaque (RM), pig-tailed macaque (PM), and human and B) among cynomolgus macaque (CM), rhesus macaque (RM), pig-tailed macaque (PM) and human.

**Supplementary Figure S3.**
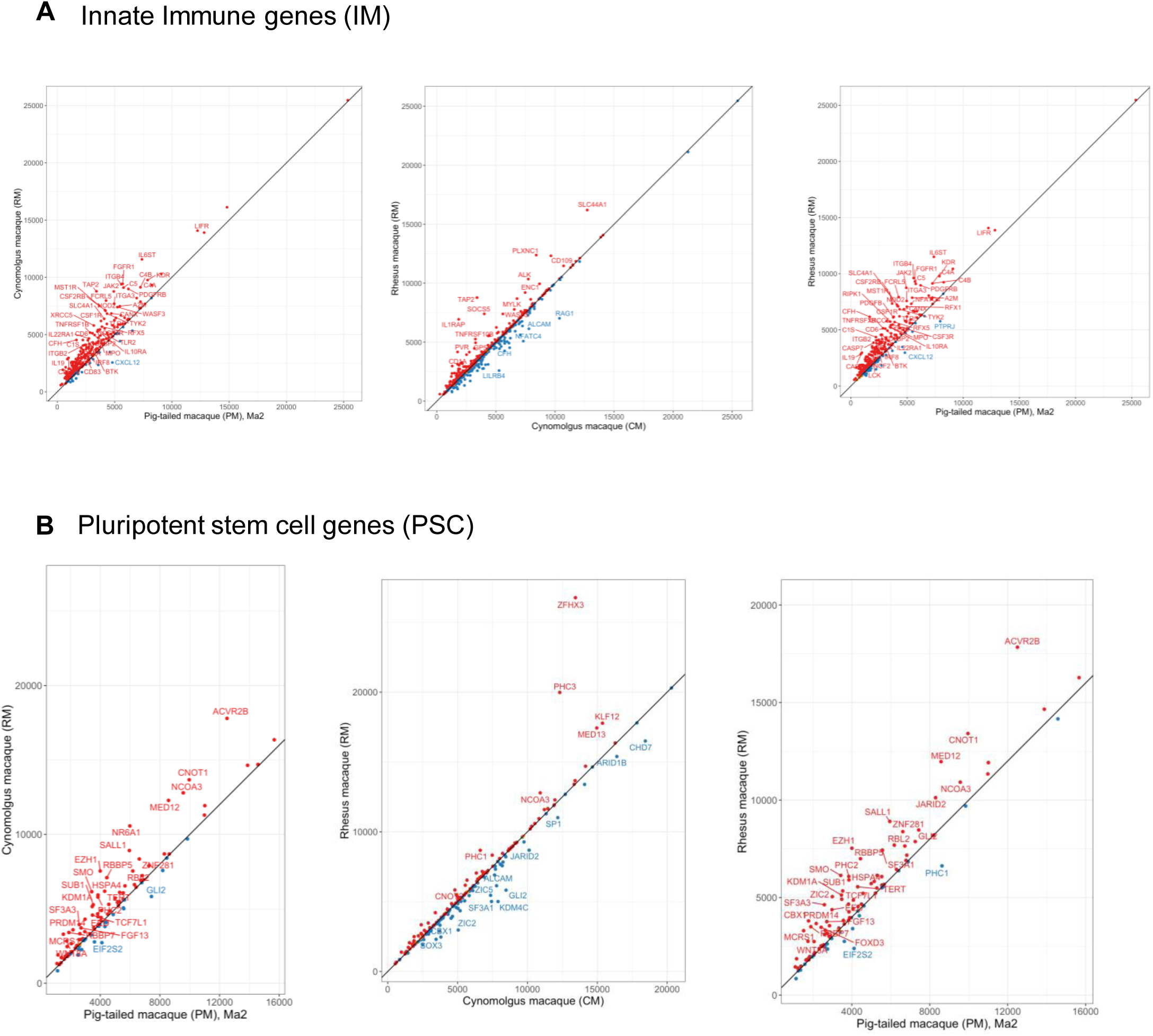
Mean bit score ortholog difference to the respective corresponding human ortholog for A) Innate immune genes (IM) between pig-tailed macaque (PM) and cynomolgus macaque (CM) (left), between cynomolgus macaque (CM) and rhesus macaque (Middle), and between pig-tailed macaque and rhesus macaque (right). The bit score differences for most of the genes are almost equal to zero. Rhesus and cynomolgus macaques, exhibited by data points clustering along the 45 degree line. B) Pluripotent stem cell genes (PSC) between pig-tailed macaque (PM) and cynomolgus macaque (CM) (left), between cynomolgus macaque (CM) and rhesus macaque (Middle), and between pig-tailed macaque and rhesus macaque (right). The bit score differences for most of the genes are almost equal to zero. Rhesus and cynomolgus macaques, exhibit data points clustering along the 45 degree line.

**Supplementary Figure S4.**
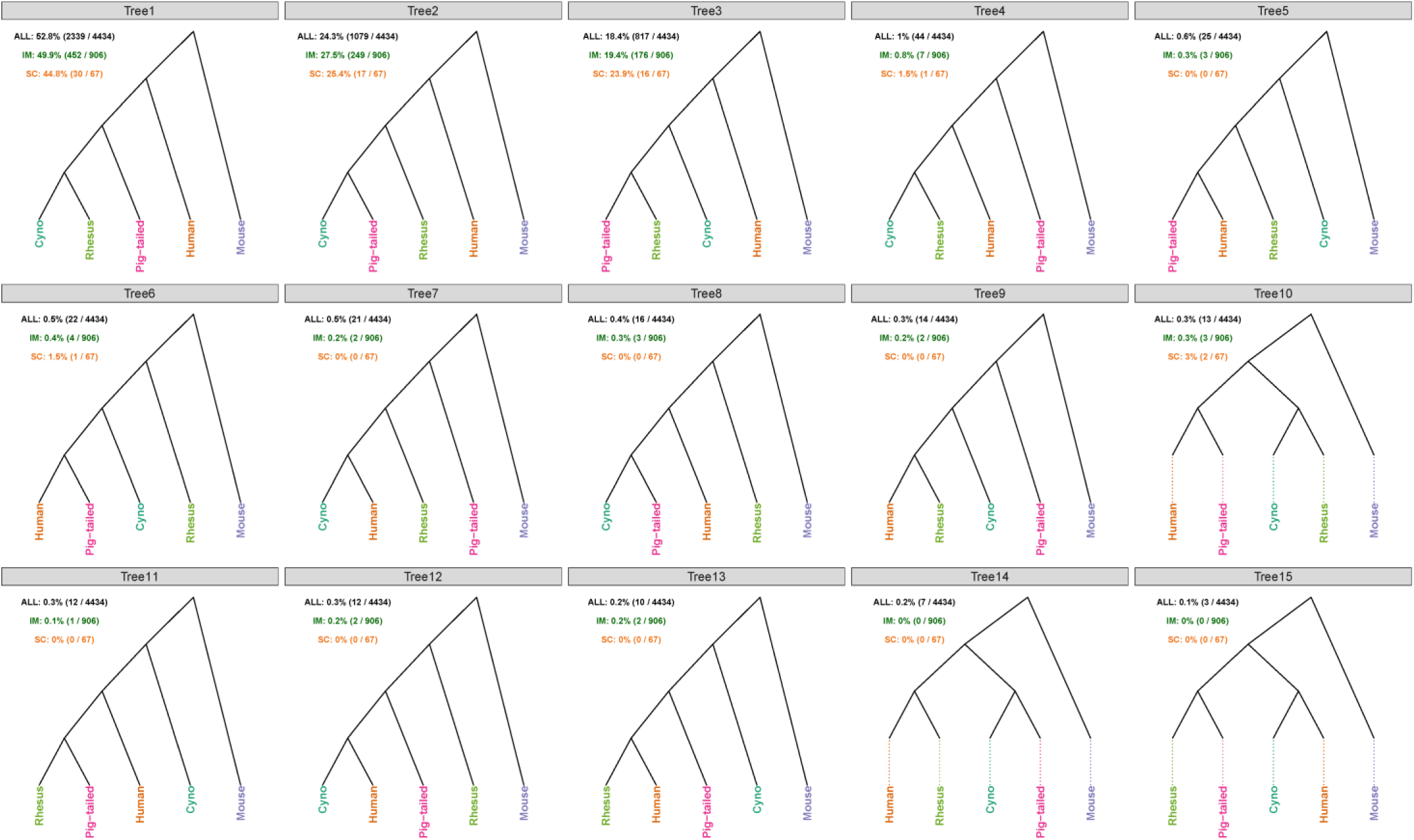
Possible phylogenetic tree topologies among three macaque species and human, using mouse as an outgroup. Tree 1 is supported by the majority of the conserved genes among the five species, 52.8% (2339/4434). Tree 6 and Tree 10 indicate a closer relationship of pig-tailed macaque to human, than either of the other two macaque species. 22 genes follow Tree 6 topology and 13 genes that follow Tree 10 topology.

